# TYPE THREE SECRETION SYSTEM-INDUCED MECHANOPORATION OF VACUOLAR MEMBRANES

**DOI:** 10.1101/2024.09.19.613840

**Authors:** Léa Swistak, Marvin Albert, Camila Valenzuela, Elif Begum Gokerkucuk, François Bontems, Anastasia D. Gazi, Stéphane Tachon, Anna Sartori-Rupp, Cammie F. Lesser, Perrine Paul-Gilloteaux, Jean-Yves Tinevez, Matthijn Vos, Jost Enninga

## Abstract

Endomembrane breaching is a crucial strategy employed by intracellular pathogens enclosed within vacuoles to access the nutrient-rich cytosol for intracellular replication. While bacteria use various mechanisms to compromise host membranes, the specific processes and factors involved are often unknown. *Shigella flexneri,* a major human pathogen, accesses the cytosol relying on the Type Three Secretion System (T3SS) and secreted effectors. Using in-cell correlative light and electron microscopy, we tracked the sequential steps of *Shigella* host cell entry. Moreover, we captured the T3SS, which projects a needle from the bacterial surface, in the process of puncturing holes in the vacuolar membrane. This initial puncture ensures disruption of the vacuolar membrane. Together this introduces the concept of mechanoporation via a bacterial secretion system as a crucial process for bacterial pathogen-induced membrane damage.

**Graphical Abstract:** 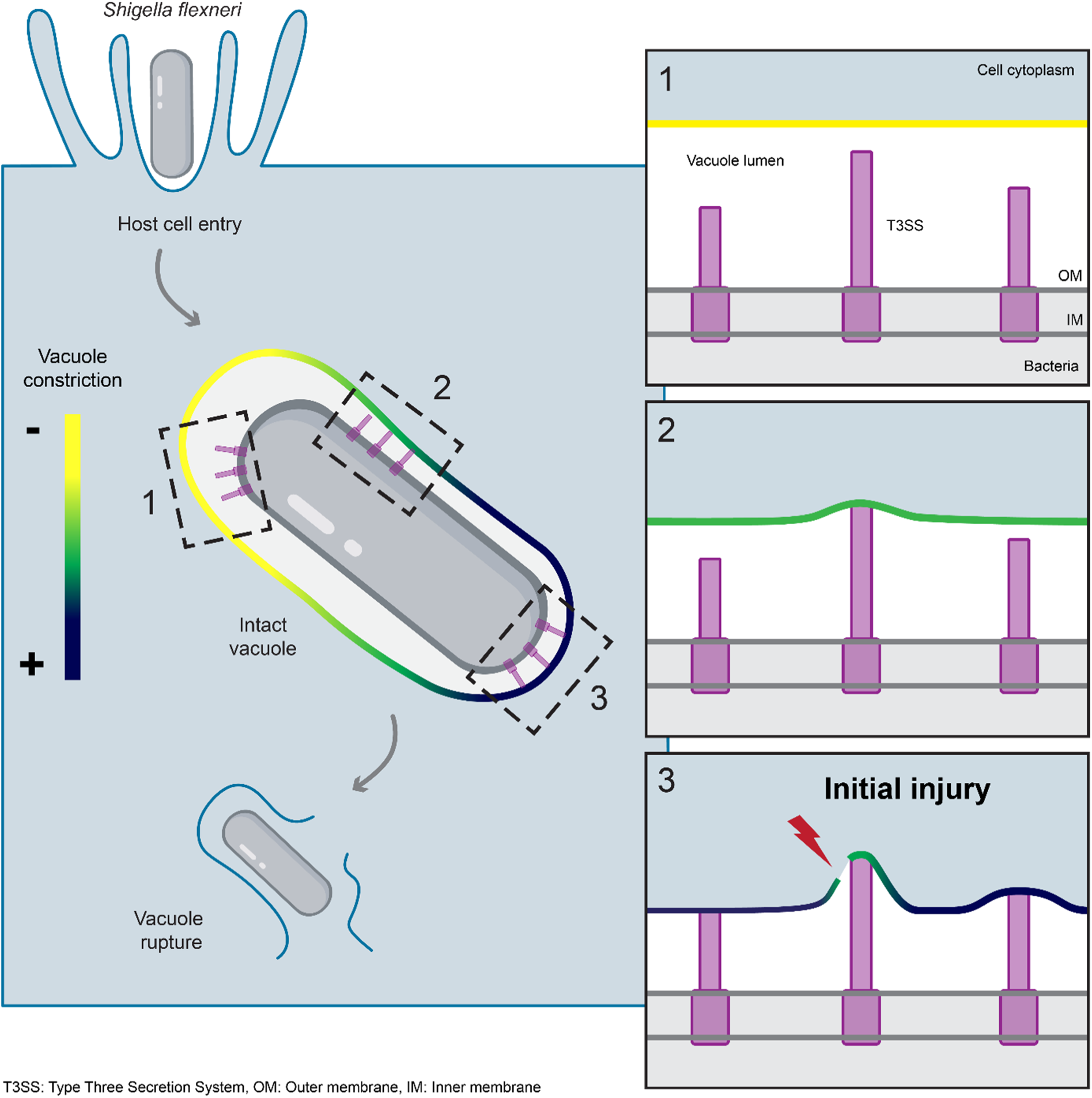

## Introduction

Deciphering the strategies of eukaryotic cell invasion by bacterial pathogens is critical to determine how they can be eliminated by the host or by antimicrobial therapies. Intracellular bacteria initially enter eukaryotic cells within vacuoles derived from the plasma membrane. Some pathogens remain vacuole-bound, blocking the trafficking towards lysosomes to avoid degradation and immune detection^1^. Other bacteria breach the vacuolar membrane to access the cytosol where they replicate and spread to neighbouring cells^2,3^. Bacterial factors such as pore-forming toxins^4^, lipids^5^, secretion systems and/or secreted effectors^6–8^, have been associated with membrane injury but the exact underlying molecular determinants or mechanisms are often unclear^9^. *Shigella flexneri* (*Shigella*) efficiently escapes its vacuole^10^, as broken vacuolar membrane remnants are actively removed from the invading bacteria^11–13^. Cytosolic access by *Shigella* relies on the Type Three Secretion System (T3SS)^14^, a specialised molecular apparatus that forms a channel, bridging the bacterial envelopes and the host cell membrane^15^. The T3SS features a basal body anchored within the bacterial envelope and a protruding needle-like structure to reach target cellular membranes for translocon pore insertion. This enables the injection of effectors proteins in one step into the host cell cytosol^15^. Initial vacuole permeabilization has been thought to be achieved via the translocon proteins IpaB/IpaC^16^, although such a mechanism has remained puzzling as many non-endomembrane damaging bacteria have T3SSs and homologous translocon complexes^15^. In particular, the dynamics from host cell entry to membrane injury and how T3SS affects vacuole integrity are unknown.

## Results

### *Shigella* T3SS drives multistep cytosolic access

We aimed to analyse the implication of the T3SS in the early steps of *Shigella* cytosolic access at high spatiotemporal resolution. For this purpose, we established a double reporter HeLa cell line expressing eGFP-Lysenin and mOrange-Galectin-3 to simultaneously track pathogen-induced vacuolar damage, and subsequent vacuolar rupture using time-lapse confocal imaging (Figure 1A). Upon vacuolar damage, sphingomyelins translocate to the cytosolic membrane leaflet, triggering detection by Lysenin^17^, which is followed by irreversible rupture of the vacuolar membrane, exposing glycans to the cytosol and allowing Galectin-3 to diffuse into the vacuolar lumen^18^. This sequence of events precedes later stages of vacuolar disassembly. We used our dual reporter to investigate the direct role of the *Shigella* T3SS in membrane breaching and elucidate its implication in vacuolar damage or rupture. For this purpose, we employed an engineered *E. coli* strain expressing functional *Shigella* T3SSs^14,19^. At 2 hours post infection, immunofluorescence showed that *E. coli* mT3*_Shigella_*^14^, invaded our reporter cell line (Figure S1). Using time-lapse microscopy, we compared the dynamics of Lysenin and Galectin-3 recruitment at the vacuole containing *Shigella* and *E. coli* mT3*_Shigella_* (Figure 1B and C). Shortly after epithelial cell entry (average 4 ± 3 min), *Shigella* vacuoles were uniformly Lysenin-positive, indicating membrane damage. This step was always followed by progressive Galectin-3 recruitment (average 7 ± 4 min after entry), consistent with the kinetics of Lysenin and Galectin-8 recruitment at the vacuole^17^, thus validating our double reporter (Figure 1D). Monitoring *E. coli* mT3*_Shigella_* cytosolic access, we noted extended delays of Galectin-3 recruitment after Lysenin signal onset (Figure 1E). In cells infected with *E. coli* mT3*_Shigella_*, the membrane remained around the bacteria even after vacuolar rupture. Such a defect in vacuole disassembly can be explained by the absence of specific effectors involved in membrane unpeeling^12^.

**Figure 1:**
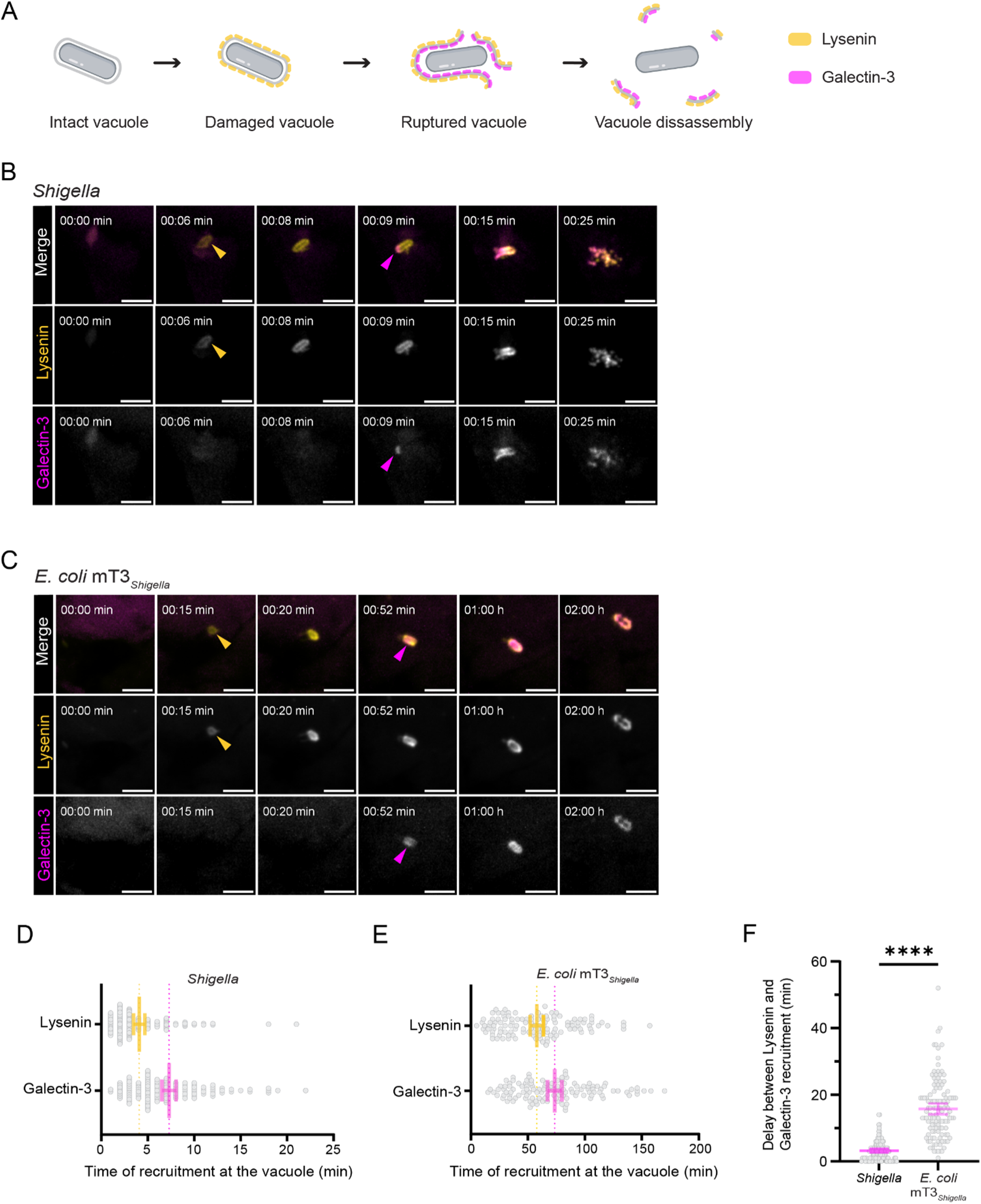
Vacuolar rupture is a multistep process that requires the T3SS for its initiation. (A) Graphical illustration of the successive steps leading to *Shigella* cytosolic access monitored using a double reporter cell line: HeLa eGFP-Lysenin mOrange-Galectin-3. Lysenin is recruited at the vacuole upon membrane damage and is followed by membrane rupture, characterized by Galectin-3 recruitment. (B and C) Representative frames (maximum intensity z-projections) of live-imaging of HeLa cells stably expressing eGFP-Lysenin and mOrange-Galectin-3 infected with *Shigella* (or *E. coli* mT3*_Shigella_* (see also Supplementary Video S1). A diffuse (B) or weak (C) signal is visible at timepoint 0 in both channels, typical for the onset of plasma membrane ruffling. Arrowheads indicate the sharp recruitment of Lysenin (yellow) or Galectin-3 (magenta). Scale bars are 5 µm. (D and E) Quantification of time of recruitment of Lysenin and Galectin-3at the *Shigella* (d) or *E. coli* mT3*_Shigella_* € vacuole after entry (time 0). *Shigella*: Lysenin, 4 ± 3 min and Galectin-3, 7 ± 4 min. *E. coli* mT3*_Shigella_*: Lysenin, 58 ± 32 min and Galectin-3, 74 ± 34 min. Bars represent the mean with 95% CI of n=120 of N=3. (F) Quantifications of the time delay between the initial recruitment of Lysenin (damage) and Galectin-3 (rupture) at the *Shigella* and *E. coli* mT3*_Shigella_* vacuoles. Mean delay of 3 ± 3 min for *Shigella* and 16 ± 9 min for *E. coli* mT3*_Shigella_*. Bars represent the mean with 95% CI, **** p <0.0001, Welch’s *t*-test. *Shigella* n=120, *E. coli* mT3*_Shigella_* n=120 of N=3.

By examining the delay between vacuolar damage and rupture through Lysenin and Galectin-3 recruitment, we found that for *Shigella*, Galectin-3 was recruited on average 3 ± 3 min after Lysenin (Figure 1F). In contrast, we observed a 5-fold delay in Galectin-3 recruitment at *E. coli* mT3*_Shigella_* vacuoles, with rupture being detected on average 16 ± 9 min after Lysenin. These results indicate that cytosolic access by *Shigella* occurs in sequential steps: vacuolar damage followed by rupture, each with distinct kinetics. This hints towards an important role of the T3SS in initiating membrane injury.

### In-cell analysis of T3SS organisation

We reasoned that close interactions between T3SSs and the vacuole would be critical to promote host endomembrane injury. We turned to in-cell cryo-electron tomography (cryo-ET), a technique allowing the visualisation of bacterial molecular machineries within infected cells^20^. To follow the involvement of the T3SS in the steps leading to vacuolar rupture, we capitalised on cryo-correlative light and electron cryo-microscopy (cryo-CLEM), imaging *Shigella*-infected HeLa cells with our stage-specific double fluorescent reporter to precisely target individual invasion steps. Vitrified cells were imaged using cryo-fluorescence microscopy (cryo-fLM) to localise fluorescently tagged *Shigella*. Guided by the cryo-fLM information, cells were thinned into lamellae at infection sites using cryo-focused ion beam (cryo-FIB) milling^21^. Zones of lamellae with intracellular bacteria were imaged by cryo-ET. After acquisition, data from all imaging modalities were correlated to gain insight into the individual bacterium invasion stage (Figure S2A-C). This cryo-CLEM approach enabled consistent data collection of intracellular *Shigella* and precise assessment of vacuolar integrity around each entering bacterium.

To examine the early stage of *Shigella* cytosolic access, we imaged cells after short infection times (10 min post-infection). At this time point bacteria were predominantly found entrapped in intact vacuoles, negative for both Lysenin and Galectin-3 (Figure 2A). We also captured the transient stage of vacuole damage, with vacuoles positive for Lysenin but negative for Galectin-3 (Figure 2B). At both stages, the vacuole tightly surrounded the bacterium, consistent with previously reported volume-imaging of *Shigella* before rupture^11^. Of note, in most of our tomograms we could spot spherical electron-densities within the tight vacuolar lumen possibly resembling small bacterial outer membrane vesicles ^22,23^ (Figure 2B, insets 1 and 2). At these infection times, the membranes from ruptured vacuoles, positive for both Lysenin and Galectin-3, were still in proximity to the invading bacteria but displayed pronounced morphological differences (Figure 2C). Indeed, it was reported that remnants of *Shigella* ruptured vacuole can either disassemble as entire segments, small pieces or fragments, or large sections may remain associated with the bacterium^12,13,24^. In accordance, segmentation of vacuolar membranes from *Shigella* exposed to the cytosol showed significant disruptions including interruptions of varying sizes, delamination, and fragmented membrane remnants (Figure 2C segmentation). Notably, damaged and ruptured vacuoles were coated by electron-dense layers that may reflect accumulation of our fluorescently tagged markers Lysenin and Galectin-3 (Figure S2D).

**Figure 2:**
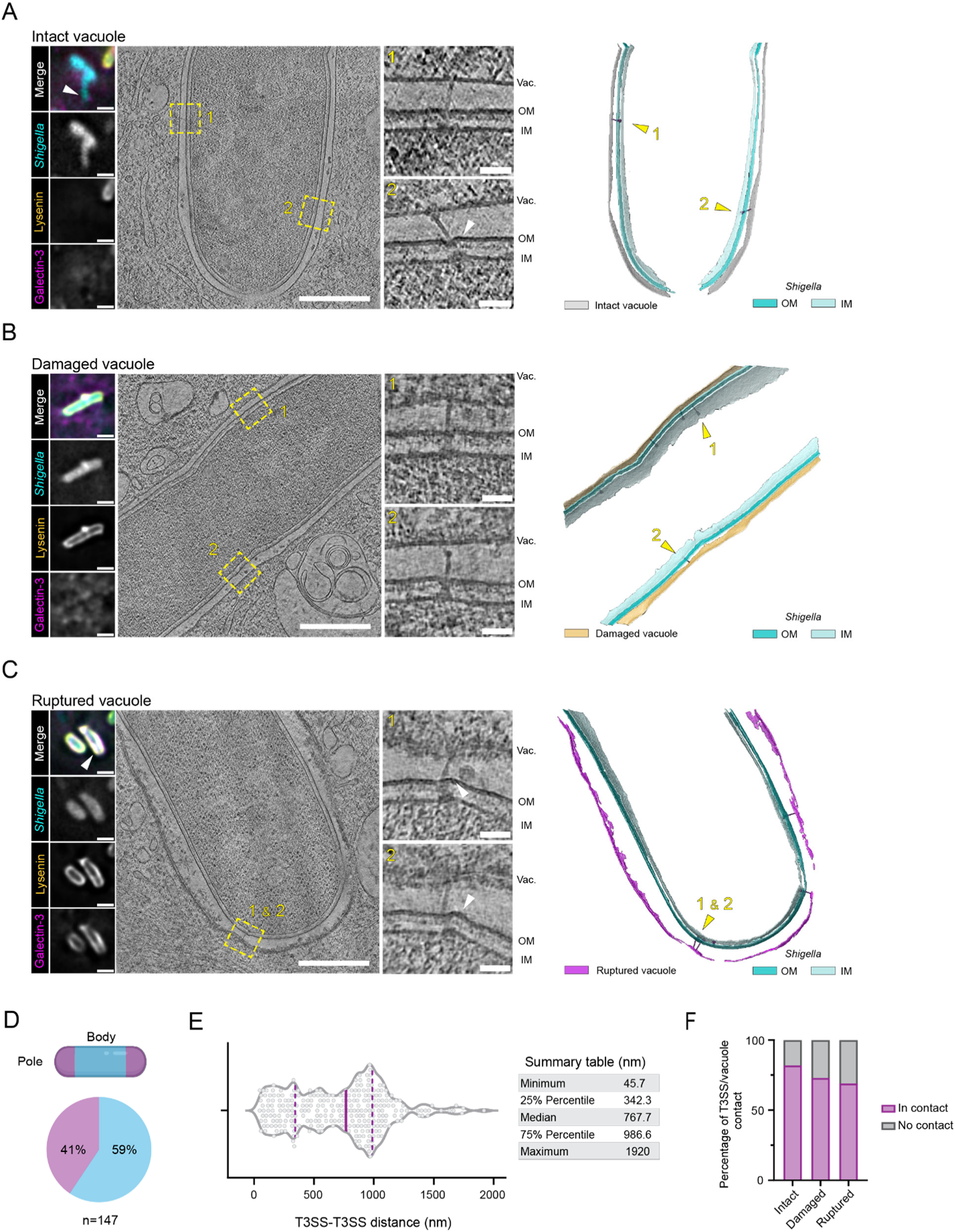
*Shigella* T3SS establishes direct contact with the vacuole membrane during infection. (A-C) HeLa eGFP-Lysenin mOrange-Galectin-3 infected with Tag-BFP *Shigella* were processed through a correlative cryo-ET workflow to capture intact (A), damaged (B), or ruptured (C) vacuoles. The first panel displays cryo-fluorescence images of the bacteria imaged by cryo-ET. Scale bars 2 µm. Second panel shows tomographic slices, with T3SSs regions marked by dashed boxes. Scale bars 500 nm. Insets show enlarged views of the T3SSs. Scale bars 50 nm. Right side panel shows membrane segmentation with vacuole coloured according to the infection stage identified by correlative cryo-ET. Yellow arrowheads indicate the T3SSs shown in the insets. White arrowheads point to bacterial membranes deformations. Vac.: Vacuole, OM: Outer membrane, IM: Inner membrane. See also Figure S2A-C for full correlation strategy and Supplementary Video S2 for tomogram and 3D rendering of vacuole and bacteria membranes. (D) Quantification of T3SSs surface distribution (pole/body) across all infection stages. Dashed lines correspond to 25% and 75% percentiles, solid line indicates the median. n=147. (E) Violin plot of the point-based calculation of distances between T3SSs across all infection stages and corresponding summary table. n=147 individual T3SS analysed. (F) Contingency graph of the percentage of T3SSs contacting or not the vacuole membrane according to the infection stage. Only the T3SSs for which a basal body and a needle were identified are plotted. Intact n=49, Damaged n=15, Ruptured n=36.

Throughout the sequential infection stages, we imaged a fair number of densities with the shape and size matching *Shigella* T3SSs (n=122) for which we distinctively identified the major substructures: the bacterial membrane spanning basal bodies and protruding needles. We also found putative basal bodies without needles (n=25, Figure S3A), suggesting that not all T3SSs are fully formed prior to bacterial entry. When T3SSs needle complexes and putative basal bodies were identified, they were frequently present in clusters. To gain general insights into the organisation of T3SSs at the bacterial surface, we assessed their distribution and found that T3SSs were located to both the poles and the bodies of the bacterial surfaces (41% placed on homogeneously curved membranes and 59% placed on straight membrane segments respectively; Figure 2D). This observation contrasts with previous reports on the polar secretion of IpaC at host cell invasion^25^ and questions a potential redistribution of the *Shigella* T3SSs from a restricted zone to the entire bacterial surface upon host entry, as observed for the T3SSs of *Chlamydia* elementary bodies^26^. To further characterise the localisation distribution of T3SSs, we annotated each T3SS, and measured the cartesian distances between nearest neighbours of individual T3SSs. While the obtained median was 767.7 nm, a subgroup (25% of total T3SSs) had shorter inter-distances, separated on average by only 195 ± 92.5 nm (Figure 2E), which agrees with our observation that T3SSs were frequently present in clusters.

T3SSs displayed a range of morphological features in relation to the vacuole membrane, either contacting it or not. Notably, some unengaged T3SSs exhibited a bulb-like density at the needle tip (Figure 2B, inset 2), which we hypothesise to be the needle tip complex, described to be larger than the needle filament of the *Salmonella* SPI-1 T3SS^27^. Nonetheless, T3SSs were mainly found associated with the vacuole during the early stages of cytosolic access (intact: 82%, damaged: 73%) and remained mostly engaged to vacuolar membrane remnants after vacuolar rupture (rupture: 69%; Figure 2F). These T3SSs displayed significant bending, with the entire structure tilted or with needles deviating from the basal body axis (Figure 2C). Previous work has shown that *Shigella* reroutes the dynein motor complex to generate forces to efficiently remove vacuolar membrane remnants from the bacteria^12^. We speculate that the observed T3SSs distortions may be induced by the forces generated during this process, which would in turn also pull on T3SSs connected to vacuolar remnants. This interpretation was further supported by deformed bacterial membranes observed around tilted T3SSs (Figure 2C, arrowheads). Notably, despite this marked phenotype suggesting the presence of mechanical forces at the T3SS-vacuole interface, bacteria with distant vacuolar membrane remnants displayed seemingly intact T3SSs at their surface with needles exposed to the host cytosol (Figure S3B). Possibly these T3SSs can be recognised by the inflammasome^28^ and may be important for priming *Shigella* secondary infections^29^.

### Vacuole tightness constrains the T3SS needle complexes

Surprisingly, T3SSs with tilted morphologies were also detected at intact vacuoles indicating that the T3SS/vacuole interface is already under substantial tension before membrane rupture. To assess the potential impact of vacuolar compression on individual T3SSs, we measured the space between vacuoles and the bacteria. During early invasion stages, the relative distance between the vacuole and the bacterial outer membrane showed tight spacing (Figure 3A) in line with the skewed distribution of calculated nearest neighbour distances (Figure 3B, grey and yellow). As expected, the space between the bacterium and the ruptured vacuole was enlarged (Figure 3A) with a wider spread distribution of vacuole-to-bacteria outer membrane nearest neighbour distances (Figure 3B, magenta). Vacuolar constriction did not change significantly from intact (48.0 ± 12.7 nm) to damaged vacuoles (48.0 ± 8.7 nm), indicating that membrane injury events occur in localised zones. The increased vacuole membrane to bacteria distances calculated for ruptured vacuoles (78.0 ± 9.2 nm) reflected general alteration of membrane integrity (Figure 3C). We then measured the needle lengths of T3SSs before cell entry (extracellular: 47.8 ± 14.4 nm) and for intracellular bacteria until vacuole rupture (vacuoles intact: 47.8 ± 10.6 nm, damaged: 49.0 ± 13.0 nm, ruptured: 48.3 ± 11.2 nm). The mean T3SS needle length was constant throughout the infection stages and matched the available luminal space before rupture, yet their measured lengths were longer and more heterogeneous than previously reported^30^. Needles that did not establish contact with the vacuolar membrane were significantly shorter, averaging 42.8 ± 13.4 nm length. Notably, T3SS needles in contact with the host membrane averaged 50.0 ± 9.6 nm, with some extending over 50 nm, exceeding the mean gap between the bacteria and the vacuole (Figure 3E). This suggests that T3SSs mediate contacts between bacteria and the vacuole and might be the space-limiting factor for vacuolar tightness.

**Figure 3:**
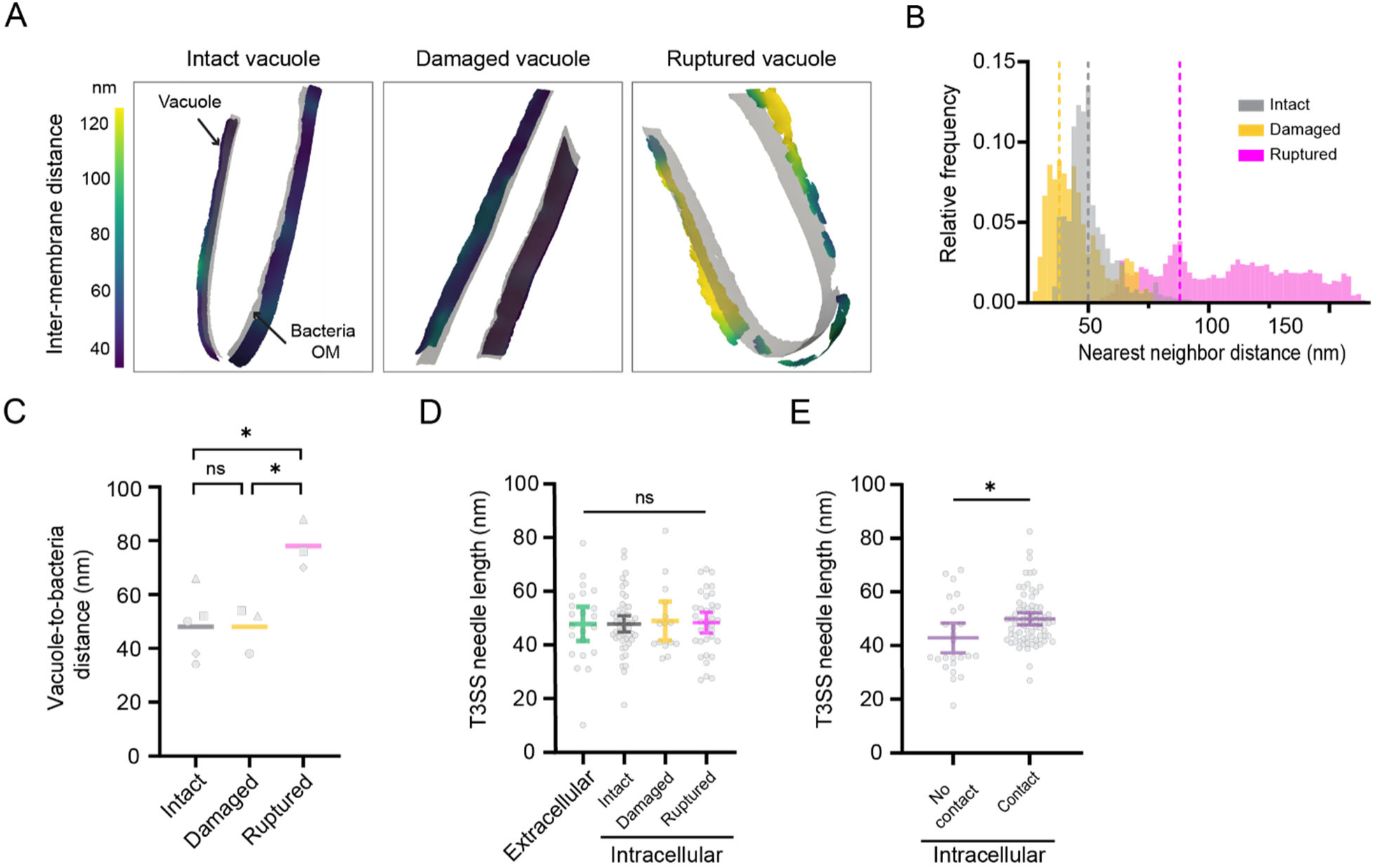
Tight vacuoles with reduced luminal space constrain the T3SS needle complexes. (A) Representative membrane surface reconstruction of intact, damaged and ruptured vacuoles coloured by 3D relative distance to the bacteria outer membrane (light grey). Arrows point to relaxed zones. OM: Outer membrane. (B) Interleaved histograms of the vacuole-to-bacteria outer membrane nearest neighbour distances of the intact (grey), damaged (yellow) and ruptured (magenta) vacuoles shown in (A). The dashed vertical lines correspond to histogram mode values. (C) Comparison of the vacuole-to-bacteria outer membrane distances at different infection stages. Histogram mode values are plotted for intact (N=5), damaged (N=3), and ruptured (N=3) vacuoles. Bars represent the mean (intact: 48.0 ± 12.7 nm, damaged: 48.0 ± 8.7 nm, ruptured: 78.0 ± 9.2 nm), * p<0.05 and ns= non-significant, one-way ANOVA with Tukey’s multiple comparisons test. (D) Quantification of the *Shigella* T3SS needle length along infection stages. Extracellular: n=22 and Intracellular: Intact n=49, Damaged n=15, Ruptured n=39. Bars represent the mean with 95% CI, ns= non-significant, one-way ANOVA with Tukey’s multiple comparisons test. (E) Quantification of *Shigella* T3SS needle length depending on whether the needles establish contact with the vacuole or not. Bars represent the mean with 95% CI, * p<0.05, Welch’s *t*-test. No contact n=25, Contact n=74.

### T3SS mechanoporation of the vacuole

At the early infection stages (Galectin-3 negative) the vacuolar membrane did not exhibit zones that were prominently perturbed. Therefore, we reasoned that early membrane injury events should be assessed at the sites where individual T3SS are in contact with the vacuole. We closely examined the T3SS-vacuole interface prior to vacuolar rupture and correlated the needle length of individual T3SS to the surrounding local vacuole-to-bacteria distances. T3SSs not contacting the vacuole had either short needles or were located to zones with relaxed vacuolar membranes (Figure S4A and B), indicating that contact requires matching between both the needle length and the inter-membrane space. This interpretation is further supported by the T3SSs with needles lengths fitting the luminal space that showed limited to no constraints on the vacuole or bacterial membranes (Figure S4C). Together, these observations imply an interplay between the needle length and the local vacuolar tightness exerting opposing constraints at the T3SS-vacuole interface.

We next focused on T3SSs with longer needles that did not fit into the tight vacuoles. T3SSs with bent insertions across bacterial membranes (Figure 4A, Figure S4D) only slightly deformed the vacuolar membrane. This indicates that the local vacuole constriction exerts a strong mechanical force on T3SSs with long needles, pushing the T3SS sideways resulting in a tilted complex. At the bacterial level, the T3SS tilt was echoed by deformations of the membranes surrounding the basal body (Figure 4A, Figure S4D arrowhead). In these cases, the local vacuole tightness is the main factor constricting the T3SSs. In other cases, T3SSs with long needles were inserted straight across the bacterial envelope and deformed the vacuolar membrane locally forming bulges. Such deformations were not detected away from the T3SS (Figure 4B), highlighting the reciprocity between needle length and vacuole constriction leading to local membrane deformation. Further examination of the vacuolar membrane morphologies at these zones revealed membrane lesions very close to the longer needle contact sites (Figure 4C and D). These T3SSs were also bent, highlighting the tensions generated by opposing forces necessary to induce membrane injuries. By segmenting the bacterial and host membranes and docking an available atomic model for the T3SS into our tomogram density we could visualise that the T3SS needle physically injured the vacuole membrane. We observed large holes localised near the long T3SS needles (Figure 4C, Supplementary Video S4) and punctures (Figure 4D). We could also spot multiple T3SSs contacting the vacuolar membrane in the same zone (Figure 4D). Even though these membrane injuries were observed, the bacteria were still negative for the vacuole rupture marker Galectin-3 (Figure S4 correlations). Thus, we propose that the *Shigella* T3SS needle complex initiates vacuole membrane injury by physically deforming it and puncturing a hole, akin to a mechanoporation^31^.

**Figure 4:**
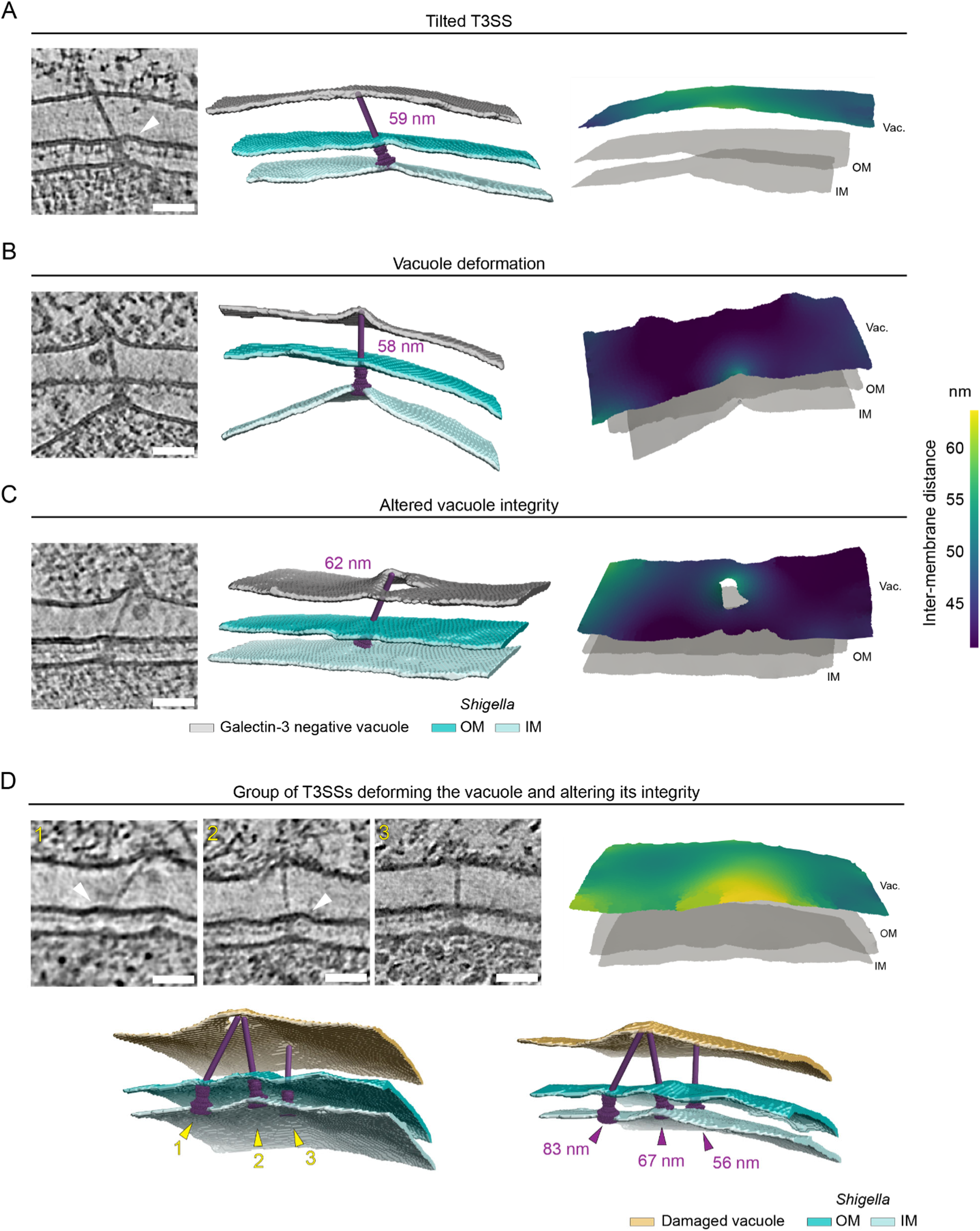
*Shigella* T3SS mechanoporation of constricted vacuolar membranes. Tomogram slices of the T3SS/vacuole zones before *Shigella* membrane rupture. White arrowheads point to bacterial membranes deformations. Scale bars 50nm. Corresponding 3D rendering of vacuole and bacteria membranes with *Shigella* T3SS 3D map (EMD-15700) fitted. Galectin-3 negative vacuole (grey), Lysenin-positive damaged vacuoles (yellow), bacteria OM (dark blue) and IM (light blue). Representative membrane surface reconstruction of the vacuole membrane coloured by 3D relative distance to the bacteria outer membrane. Bacteria membranes are shown in light grey. Vac.: Vacuole, OM: Outer membrane, IM: Inner membrane. See also Figure S4E-G for correlations and T3SS positioning information and associated Supplementary Video S3. (A) T3SS with needle length (59 nm) exceeding local luminal space (∼50 nm) adopting a tilted insertion within bacteria membranes with OM showing deformation. The vacuole membrane is only slightly deformed at the T3SS contact site. (B) Straight T3SS with needle length (58 nm) exceeding local luminal space (∼45 nm), pushing the vacuole membrane forming a steep bulge at T3SS/vacuole contact site. (C) T3SS with needle length (62 nm) exceeding local luminal space (∼45 nm) deforming the vacuole membrane up to altering its integrity by physically puncturing it. See also Supplementary Video S4. (D) Group of three T3SS localising in the same zone of the bacteria surface. T3SS n°1 and 2 contact, deform, and perforate the vacuole membrane at the same spot. T3SS n°1 needle (83 nm) is long and seems to deviate from basal body axis showing that strong constrains between T3SS and the vacuole at are play. The locally relaxed vacuole to bacteria space (∼50-55 nm) potentially limits additional damage by the T3SS n°3 needle (56 nm) that only slightly deforms the membrane.

## Discussion

We previously described *Shigella* vacuolar escape^12^, but the mechanism of initial vacuolar membrane injury has remained unclear. Here, we address this knowledge gap showing how the T3SS breaches the vacuolar membrane, during the first step of cytosolic access. Our data indicate that the T3SS/vacuole interface is under major tension before vacuole rupture, with opposing forces from T3SS needle complexes and vacuolar constriction. Membrane injury occurs in the vicinity of T3SSs that do not fit the tight vacuolar lumen. *Shigella* T3SSs contact host membranes at cholesterol-rich nanodomains^32^, with increased stiffness^33^. In case of limited space, these sites appear to be prone to T3SS-induced membrane injury. Moreover, the *Shigella* vacuole is enclosed in a thick actin cocoon^13^ a structure that possibly supports membrane resistance to deformations induced by the T3SS needles. The process identified here is highly reminiscent to membrane injury due to mechanoporation. Together, the tight *Shigella* vacuole, limited lipid availability and the surrounding actin cytoskeleton might prevent spontaneous resealing of T3SS-induced holes leading to catastrophic membrane injury^34^. We argue that *Shigella* is highly efficient exerting T3SS-induced mechanoporation, but it may also happen in other T3SS-bearing pathogens, such as *Salmonella*. Accordingly, there is a correlation between the shrinking of the *Salmonella* containing vacuole^35,36^ and endomembrane breaching^37^. *Yersinia* has also been described to deform the phagosome membrane at the interface with the T3SS however injuries were not observed^38^. In line with this, other pathogens with canonically larger vacuoles do not cause endomembrane breaching ^39,40^. We cannot exclude that other pathogens could still damage the vacuole through T3SS-mediated mechanoporation but have evolved additional mechanisms of hijacking host membrane repair pathways, preventing vacuole rupture and cytosolic escape^41,42^. Furthermore mechanoporation may also take place during endomembrane damage through other secretion systems such as Type 6 Secretion System^43^. Together, we establish a new paradigm in membrane damage by pathogens with a direct involvement of bacterial secretion system during early membrane injury.

## Supporting information

Video S1. Time-lapse microscopy monitoring of Lysenin and Galectin-3 recruitment at the Shigella and E. coli mT3Shigella vacuole, related to Figure 1B

Video S2. Tomograms and corresponding segmentations of the distinct stages of Shigella cytosolic access, related to Figure 2A-C

Video S3. Tomograms and corresponding segmentations of Shigella prior cytosolic access, related to Figure S4E-G

Video S4. T3SS punctures host endomembrane, related to Figure 4C

## Acknowledgments

We thank Sandrine Schmutz (Flow Cytometry Platform) of C2RT for technical assistance during cell sorting. We also thank Mikhail Kudryashev, Patricia Bassereau, Félix Rey and Javier Pizarro-Cerda for discussions in the assembly of the manuscript and Felix Randow for sharing tools used in this study. Work in the unit of J.E. is supported by the ERC (CoG Endosubvert) and the ANR (grants HBP sensing, PureMagRupture, and RabReprogram) and is a member of the LabEx “IBEID” and “Milieu Interieur”. L.S. was supported by École Doctorale 562 BioSPC, Université de Paris-Cité and by the Fondation pour la Recherche Médicale, grant number FDT202204014967. C.F.L. has been supported by a grant from the NIAID (R01 AI128360). Furthermore, we would like to thank the Domaine d’Intérêt Majeur ELICT 2020 (DIM Elicit 2020, grant “Ultrapath”) from the Région Ile-de-France and the National Infrastructure France-BioImaging.

## Author contributions

L.S. performed infections with help of C.V. for *E. coli* mT3 strains. C.F.L. provided essential tools. E.B.G. generated and L.S. validated the double reporter cell line. L.S. collected, analysed, and interpreted all microscopy data. A.S.R. aided in cryo-fLM and cryo-ET data acquisition. S.T. participated in cryo-FIB milling and cryo-ET data acquisition. F.B. designed the tomogram processing pipeline with help of L.S. for implementation. M.A. and P.P.G design the post-acquisition correlation workflow implemented by M.A., supervised by J.Y.T with input from L.S. M.A. performed data analysis (membrane surface reconstructions and distance measurements). A.G. and C.V. aided L.S. in interpreting and visualizing the results. L.S. and J.E. wrote the manuscript with help by C.V. and input from all authors. M.V. and J.E. supervised the project. L.S. and J.E. conceptualized the study.

## Material and Methods

### Cell culture and stable cell line generation

All HeLa epithelial cell lines used in this study were cultured in Dulbecco’s modified Eagle’s medium (DMEM, Gibco) supplemented with 10% (v/v) heat-inactivated foetal bovine serum (Sigma-Aldrich) at 37°C, 5% CO_2_ and were continuously monitored for mycoplasma.

The double expressing mOrange-Galectin-3 and eGFP-Lysenin stable cell line was generated using the Sleeping Beauty (SB) transposon system^44^. Briefly, N-terminally eGFP tagged Lysenin was PCR amplified from M6P-GFP-LyseninW20A plasmid^17^ (kindly provided by Felix Randow) using the primers Forward_eGFP (5’aggcctctgaggccaccatggtgagcaagggcgag 3’) and Reverse_Lysenin (5’ aggcctgacaggcctcagcccacgacttccagga 3’). The obtained amplicon was cloned into the Sleeping Beauty transposon (pSBbi-BLA, addgene plasmid #60526) using the Sfil sites. The resulting pSBbi eGFP-LyseninW20A plasmid was confirmed by sequencing and co-transfected with the vector encoding the SB100X transposase, pCMV(CAT)T7-SB100X (Addgene plasmid #34879) in the monoclonal HeLa pSBbi mOrange Galectin-3^13^ (puromycin resistant). 48 hours post-transfection, cells were switched to selection media containing 10 µg/mL blasticidin (Gibco) and 1.7 µg/mL puromycin (Gibco). After 10 days of culturing in selection media, cells were sorted to select double fluorescence positive clones (eGFP-Lysenin and mOrange-Galectin-3) using a BD FACSAria III Cell Sorter (BD Biosciences). A polyclonal subpopulation with high expression of eGFP-Lysenin was further selected for expansion.

### Bacterial strains and infection for fluorescence microscopy

*Escherichia coli* DH10β (Thermo Scientific), derivative minimal T3SS (mT3) strains mT3 *E. coli* (pLLX13 *ipaJ* thru *spa40*, TET^R^, KAN^R^ + pNG162-VirB)^19^ and mT3Δeff_IpaA (pLLX13 *ipaJ* thru *spa40*Δ*ipgB1*,Δ*icsB*,Δ*ipgD*,TET^R^, KAN^R^ + pNG162-VirB)^14^ were grown in lysogeny broth (LB) at 37°C with shaking. For the mT3 strains, LB was supplemented with 20 µg/mL tetracycline, 50 µg/mL kanamycin, and 100 µg/mL spectinomycin (all from Sigma-Aldrich). For *E. coli* infection experiments, bacteria were diluted in LB at 1:50 from an overnight culture and grown at 37°C. In the case of the mT3 strains, T3SS expression was induced with 1mM IPTG (Thermo Scientific). At an optical density of 600 nm (OD_600_) ∼0.45 bacteria were harvested by spinning and washed twice in EM buffer (120 mM NaCl, 7 mM KCl, 1.8 mM CaCl2, 0.8 mM MgCl2, 5 mM glucose, 25 mM HEPES, pH 7.3). Bacteria were diluted to a multiplicity of infection (MOI) of 100 in EM buffer and spun down on the cells for 10 min at 180g, at room temperature (RT).

All *Shigella* strains were derivatives of the WT *Shigella flexneri* M90T, when indicated they carried the pGG2-eGFP or pGG2-TagBFP^45^ plasmids, for constitutive expression of eGFP or TagBFP, respectively. *Shigella* strains were grown in trypticase soy broth (TCS) medium at 220 rpm at 37°C, when applicable the medium was supplemented with 100 µg/mL ampicillin. On the day of the infection, bacteria were sub-cultured from an overnight culture in fresh TCS at 1:100 dilution until OD_600_ reached ∼0.45. Bacteria were harvested, washed once in EM buffer, and incubated for 15 minutes at 37°C with shaking in EM buffer supplemented with 1 µg/mL poly-L-lysine hydrobromide (Sigma-Aldrich). The bacterial solution was washed twice, and the bacteria were diluted to the appropriate MOI.

### Fluorescence microscopy and immunolabeling

Imaging was carried out on a Nikon Ti-E inverted microscope equipped with a Perfect Focus System (PFS), a spinning disk confocal system (CSU-W1, Yokogawa), and an ORCA flash 4.0 camera (Hamamatsu). Time-lapse imaging was performed using a CFI S Plan Fluor ELWD 40x/0.60 air immersion objective (MRH08430, Nikon) and microscopy of fixed samples with a CFI Plan Apo VC 60X/1.2 water immersion objective (MRD07602, Nikon). For fluorescent microscopy experiments, 40 000 cells were seeded into 8-well glass bottom microslides (ibidi). For time-lapse imaging, the microscope chamber was heated at 37°C, and bacterial invasion was monitored immediately after bacteria addition to the cells. Images were recorded every minute for 2.5 hours (*Shigella*) or 3.5 hours *E. coli* mT3*_Shigella_*. For microscopy of fixed samples, cells were infected with *E. coli* strains at 37°C, 5% CO_2_ for 2h, while *Shigella* infections were carried out for 30 minutes at 37°C. The inside-out staining protocol was adapted from another study^46^. Infected cells were washed three times in DPBS (Gibco) and fixed in 3.7% (wt/vol) Paraformaldehyde (PFA, Electron Microscopy Science), diluted in DPBS for 20 min at RT. All antibodies were diluted in an immunolabelling solution (2% BSA in DPBS) and incubated for 30 min at RT. First, the extracellular bacteria were labelled with a rabbit polyclonal antibody to *E. coli* (1:2000; ab137967 Abcam) or a rabbit polyclonal antibody to *Shigella* (1:2000; ab65282 Abcam). Cells were then washed three times with DPBS and permeabilized with 0.1% Triton X-100 in DPBS for 10 min at RT. Permeabilization solution was washed three times with DPBS. The extracellular bacteria were labelled with goat anti-rabbit AlexaFluor™ 647 (1:2000; A32733, Invitrogen), and the total bacteria were stained with DAPI (1:2000, Fisher Scientific). Finally, samples were washed three times with DPBS and stored hydrated at 4°C, protected from light. z-stacks of individual positions were recorded with a 0.3 µm step size.

### Time-laps image analysis

Images were analysed using FIJI^47^. Images from the time-lapse series were corrected for the intensity decay due to photobleaching using the FIJI bleach correction plugin^48^. Vacuole damage and rupture times were manually quantified.

### *S**higella* epithelial cell infection for cryo-electron microscopy

Cryo-EM gold grids (Quantifoil R2/2 Au 200 mesh, Quantifoil) were placed in the slots of a custom-made PDMS stencil (Alvéole) on a 35 mm dish (µ-Dish 35 mm, ibidi). The montage was plasma-cleaned for 45 seconds and sterilized under UV irradiation in a laminar flow hood for 15 min. Cell culture medium was added to the dish and equilibrated for 15 min at 37°C. 85 000-100 000 HeLa cells were seeded and grown for 24 hours at 37°C, 5% CO_2_. Before the infection, cells were washed 3 times with DMEM and incubated with bacterial suspension at a high MOI (100 to 400). Infection was synchronized at RT for 15 min and cells were infected at 37°C for 10- or 20-min. Cells were washed three times with DPBS (Gibco) and fixed for 15 min in 2% PFA (Electron Microscopy Science), 0.05% Glutaraldehyde (Sigma-Aldrich) in 0.1M HEPES (Gibco) followed by 15 min in 4% PFA, 0.1% Glutaraldehyde in 0.1M HEPES. Fixed cells were washed 3 times with DPBS and immediately vitrified. Grids were blotted from the backside and plunged into liquid ethane at liquid nitrogen temperature using a Leica EM GP automatic plunger with the following settings: humidity 98%, blot time: 8 seconds, 20°C. The vitrified grids were stored in sealed boxes in liquid nitrogen until further use.

### Fluorescence-guided cryo-FIB milling

Grids clipped into cryo-Focused ion Beam (FIB) milling Autogrids (Thermo Fisher Scientific) were imaged in light microscopy using a Leica THUNDER Imager EM Cryo-CLEM (Leica Microsystems). A global map of the grid was acquired to aid later correlations and individual z-stacks with 0.35 µm spacing were recorded at the regions of interest. Cryo-FIB lamellae were prepared using an Aquilos 2 dual-beam cryo-focused ion beam scanning electron microscope (cryo-FIB-SEM; Thermo Fisher Scientific) instrument equipped with a cryo-transfer system, a cryo-stage, and a 45° pre-tilted shuttle (Thermo Fisher Scientific). In order to locate region of interest for the lamellae milling, a SEM map of the grid was acquired and cryo-fluorescence images were loaded at the correct relative position and orientation within the SEM map using Maps 3.2 (Thermo Fisher Scientific). Before milling an organometallic platinum layer was deposited on the cells for 1 min using the gas injection system. Lamellae were automatically milled overnight to 1 µm with a reducing current (1 nA to 0.1 nA) using AutoTEM (v.2.2, Thermo Fisher Scientific) with a 10° milling angle. The next day, lamellae were manually polished to ∼200 nm and stored in liquid nitrogen.

### Tilt series data collection and reconstruction

Cryo-ET datasets were collected using SerialEM software^49^ (version 4.1.0 to 4.1.4) on a 300 kV cold-Field Emission Gun Titan Krios transmission electron microscope (Thermo Fisher Scientific) equipped with a Falcon 4i direct electron detection camera (Thermo Fisher Scientific) and Selectris X an energy filter (Thermo Fisher Scientific). For tilt series acquisition the microscope was set in nanoprobe mode, energy filter at 20 eV (zero loss) and an objective aperture of 100. Tilt series were acquired with a dose dose-symmetric acquisition scheme^50^ ranging from +70° to −50° starting from a 10° pretilt with 3° increments at the nominal magnification of 26 000x and 42 000x corresponding to a pixel size of 3.104 and 4.81 Å respectively with defocus ranging from 3 to 5 µm. The total dose applied was 140 e^-^/Å^2^ with an electron dose starting at 3.5 e^-^/Å^2^ at 10° pretilt and exposure time varying to be 1.15 times higher at high tilts. The cold-FEG was flashed before each tilt series acquisition and frames were saved in eer format. Tilt series were processed with our in-house reconstruction script. First, the frames were motion-corrected and gain-corrected using MotionCor2^51^. 2D contrast transfer function (CTF) was calculated with ctfplotter and the correction was applied with phase-flipping using ctfphaseflip^52^. Tilt series were dose-weighted and aligned in AreTomo^53^ and tomograms were reconstructed using IMOD by weighted back-projection ^54^ to a pixel size ∼10 Å corresponding to a binning of 2 or 3 depending on the dataset and applying and “exact filter”. Finally, reconstructed tomograms were filtered using isotropic reconstruction software IsoNet^55^ with the following parameters (make_mask --density_percentage 50 --std_percentage 5; refine --iterations 30 --noise_start_iter 10,15,20,25 --noise_level 0.05,0.1,0.15,0.2). Tomograms presented in the supplementary movies were denoised with Topaz-Denoise^56^.

### Tomogram analysis

#### Image correlation

Post acquisition image correlation was performed using a semi-automatic workflow. All images were placed into a world coordinate system by extracting their relative stage coordinates, pixel spacing and orientation from image metadata. First, precise spatial transformations between EM images at different resolutions was determined in an automated manner. For this, all EM montages were stitched using the python package multiview-stitcher^57^. Then, for each tomogram the map with the highest resolution containing its central coordinate was determined, i.e. a preview or anchor map. This map was then registered against its closest medium magnification map, which was subsequently registered against the low magnification map representing the grid level (using translation, rotation and uniform scaling). The so obtained transformations were then used to precisely place all EM images jointly onto the cryo-EM grid. Registration between the cryo-EM and fluorescent modalities was then performed at the grid level by estimating an affine transform from manually placed landmarks using the napari plugin affinder (https://github.com/jni/affinder). Finally, stacks containing transformed and cropped views of all relevant imaging modalities data around each tomogram position were generated at low, medium, and high resolutions and exported as composite tiff files. Python scripts related to this and other methods described in this publication are available under https://gitlab.pasteur.fr/iah/2024_swistak_et_al_code and make use of the scientific python ecosystem^58–64^. For figure preparation, channels of cryo-fLM images were registered using the FIJI Rigid Registration plugin^47^ to correct for chromatic shift.

#### T3SS annotation and measurements

IsoNet-filtered tomograms were manually annotated using IMOD 3D visualization tools (3dmod and slicer)^65^. Individual objects were created for each secretion system and points were placed in the inner membrane/basal body interface (point 1), basal body/needle junction or basal body/bacteria outer membrane junction for the basal body only configuration (point 2), and the needle tip or vacuole membrane/T3SS tip interface (point 3) in this strict order. The obtained point coordinates were loaded into python using the package imodmodel (https://github.com/teamtomo/imodmodel). Distances between the T3SS points were calculated as the Euclidean distances between the manually determined 3D coordinates. For inter-T3SS distances, point 2 was considered as reference. Substacks around T3SSs were extracted by aligning their y axis with the vector 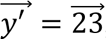, the x axis with 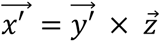 and the z axis with 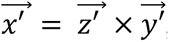, where 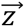 indicates the z axis of the tomogram and x the vector product.

#### Semi-automated segmentation

Membranes of full tomograms or T3SSs substacks were automatically segmented using the MemBrain-Seg^66^ segmentation tool (v2, v9b pretrained model). The resulting segmentations were imported into napari^67^ for manual refinement. Segmented objects belonging to the inner and outer membranes of bacteria and vacuole membranes were annotated and manually curated to remove false positive detections. For display purposes, curated segmentations were loaded into UCSF ChimeraX^68^ (v1.8), and *Shigella* T3SS atomic model (EMD-15700) was docked into our tomogram densities to indicate T3SS positions on bacterial surface.

#### membrane morphometrics – inter membrane distance calculation and representation

Surface meshes were reconstructed from the connected components of the curated segmentation masks using the vtkSurfaceReconstruction filter of the Visualization Toolkit^69^ within pyvista^70^. The distances between membranes were calculated by nearest-neighbour analysis. To avoid overestimating distances at surface edges, we excluded distance measurements for points with nearest neighbour located at the edges of the reference surface. Instead, for those points, we considered the same distance as was assigned to the closest point within the same surface. For visualization, mesh borders were smoothed, and 3D renderings were created using pyvista.

### Statistical analysis

Two-tailed unpaired *t*-tests and ANOVA with Tukey’s multiple comparisons tests were performed using GraphPad Prism version 10.2.2 for Windows, GraphPad Software, Boston, Massachusetts USA (www.graphpad.com). With a p value considered significant if p <0.05 with *<0.05 and ****<0.0001.

### Figure preparation

All figures were prepared using Adobe Illustrator (v. 27.8.1) and some elements of Figure 1A and S2 were created with BioRender.com.

## Data and code availability

The datasets generated during and/or analysed during the current study are available from the corresponding author J.E. on reasonable request. Codes of the methods described in this publication are available under: https://gitlab.pasteur.fr/iah-public/2024_swistak_et_al_code.

## Supplemental information

**Figure S1:**
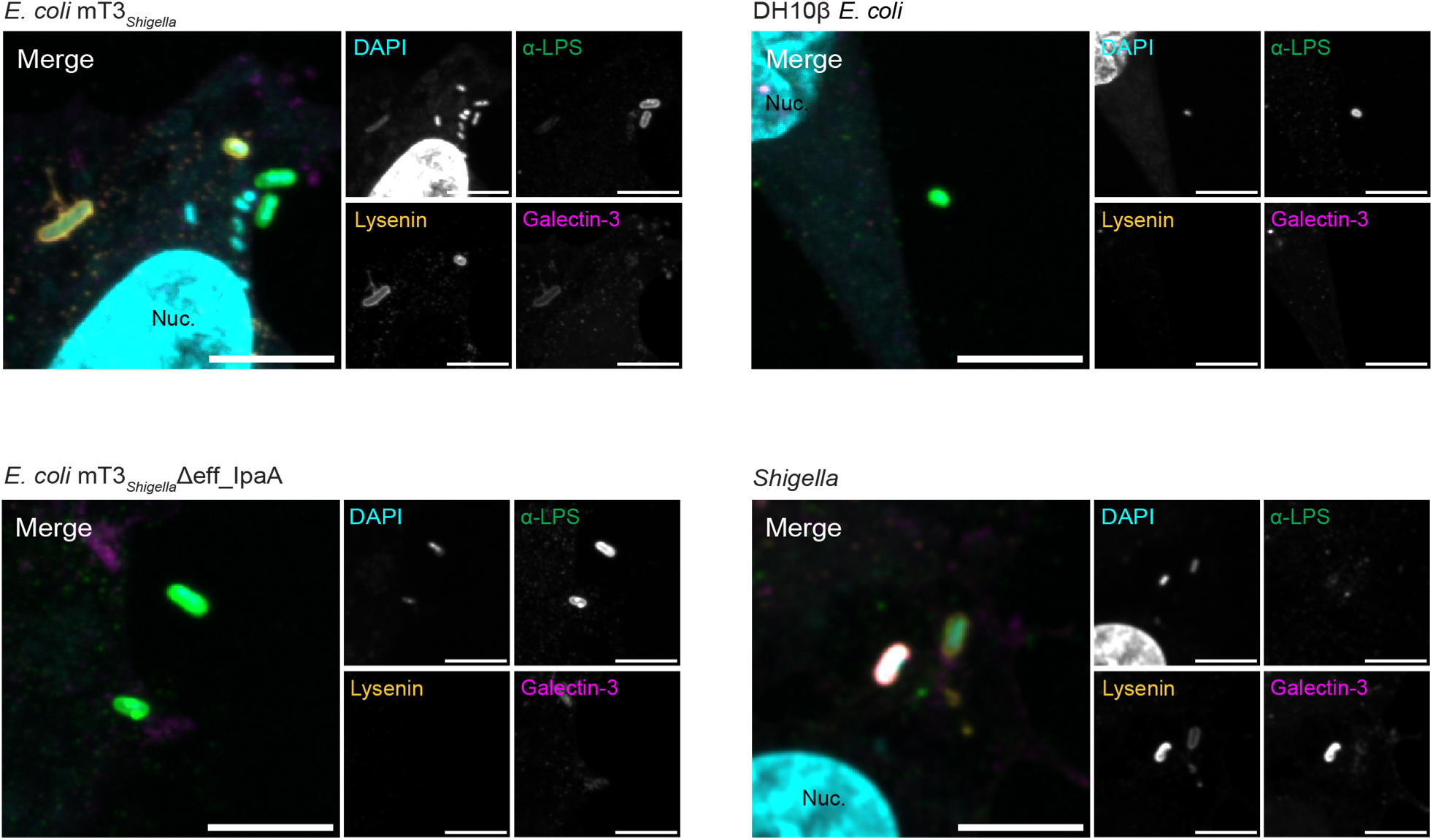
A minimal set of effectors is required for efficient *E. coli* mT3*_Shigella_* entry into epithelial cells. Representative microscopy images of HeLa eGFP-Lysenin mOrange-Galectin-3 cells infected with either *E. coli* mT3*_Shigella_*, *E. coli* mT3*_Shigella_*Δeff_IpaA, DH10β *E. coli* for 2 hours or WT *Shigella* for 30 minutes. Extracellular bacteria were stained with an antibody directed against LPS and total bacteria (extra and intracellular) were labelled with DAPI. Only Shigella and *E. coli* mT3*_Shigella_* invade HeLa cells. eGFP-Lysenin: yellow, mOrange-Galectin-3: magenta, DAPI: cyan, and α-LPS: green. Intracellular bacteria: cyan, Extracellular bacteria: cyan and green. Scale bars 8 µm.

**Figure S2:**
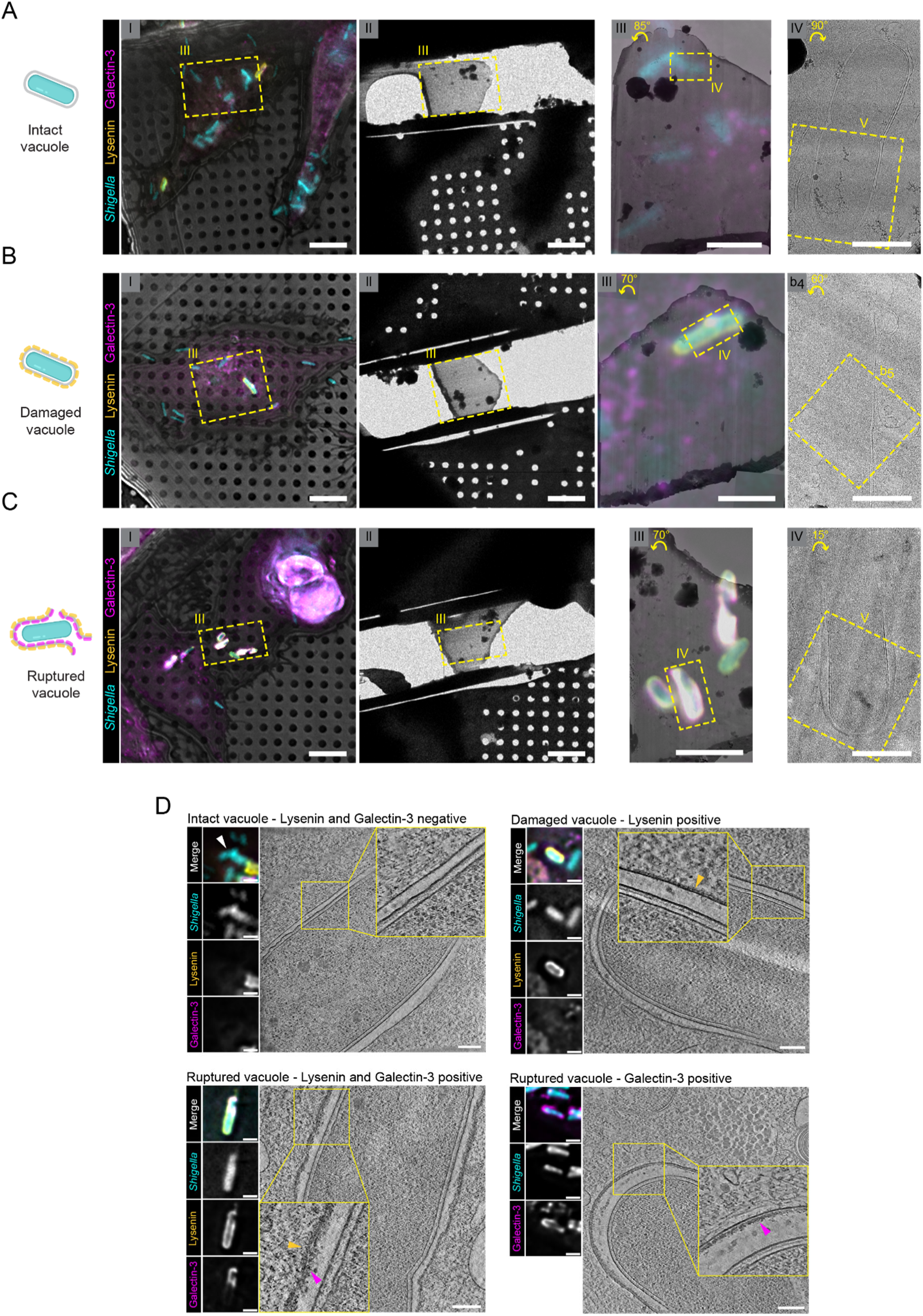
Correlative cryo-ET recapitulates the successive steps of *Shigella* vacuole damage and rupture. (A-C) Detailed correlation steps for the identification intact (A), damaged (B), or ruptured (C) vacuoles (shown in Figure 2) in HeLa eGFP-Lysenin mOrange-Galectin-3 cells infected with Tag-BFP *Shigella* processed through a correlative cryo-ET workflow.

(I) Vitrified cells on cryo-EM grids were imaged by cryo-fluorescence microscopy to localise infection sites and target them for lamella milling. Tag-BFP *Shigella*: cyan, eGFP-Lysenin: yellow, and mOrange-Galectin-3: magenta. Scale bars: 10 µm.
(II) Cryo-TEM overview of the region targeted in I after cells were thinned into lamellae using cryo-FIB-milling. Scale bars 10 µm.
(III) Cryo-lamellae maps overlayed with the corresponding cryo-fluorescence images. Scale bar 5 µm. The rotation angles from images I and II to III are indicated on the top left.
(IV) Inset of III showing the bacteria that was targeted for imaging. Scale bar 1 µm. The rotation angles from images III to IV are indicated on the top left. Square boxes V correspond to regions presented in the tomographic slices of Figure 2A, B and C respectively. (D) Slice through tomograms (scale bars 200 nm) and corresponding cryo-fLM images (scale bars 2 µm) of *Shigella* (white arrowhead) at different infections stages with vacuole membranes showing different coating patterns that may reflect on recruitment of overexpressed Lysenin (yellow arrowhead) and Galectin-3 (magenta arrowhead). Intact vacuoles: membrane coating is never observed. Damaged vacuoles: cytosolic side of the vacuole membrane uniformly coated with electron-dense layers upon Lysenin recruitment. Ruptured vacuoles: both on the cytosolic and luminal side of Lysenin and Galectin-3 double-positive vacuoles are coated while only the luminal side of the vacuole membrane is coated in cells expressing just the Galectin-3 marker.

**Figure S3:**
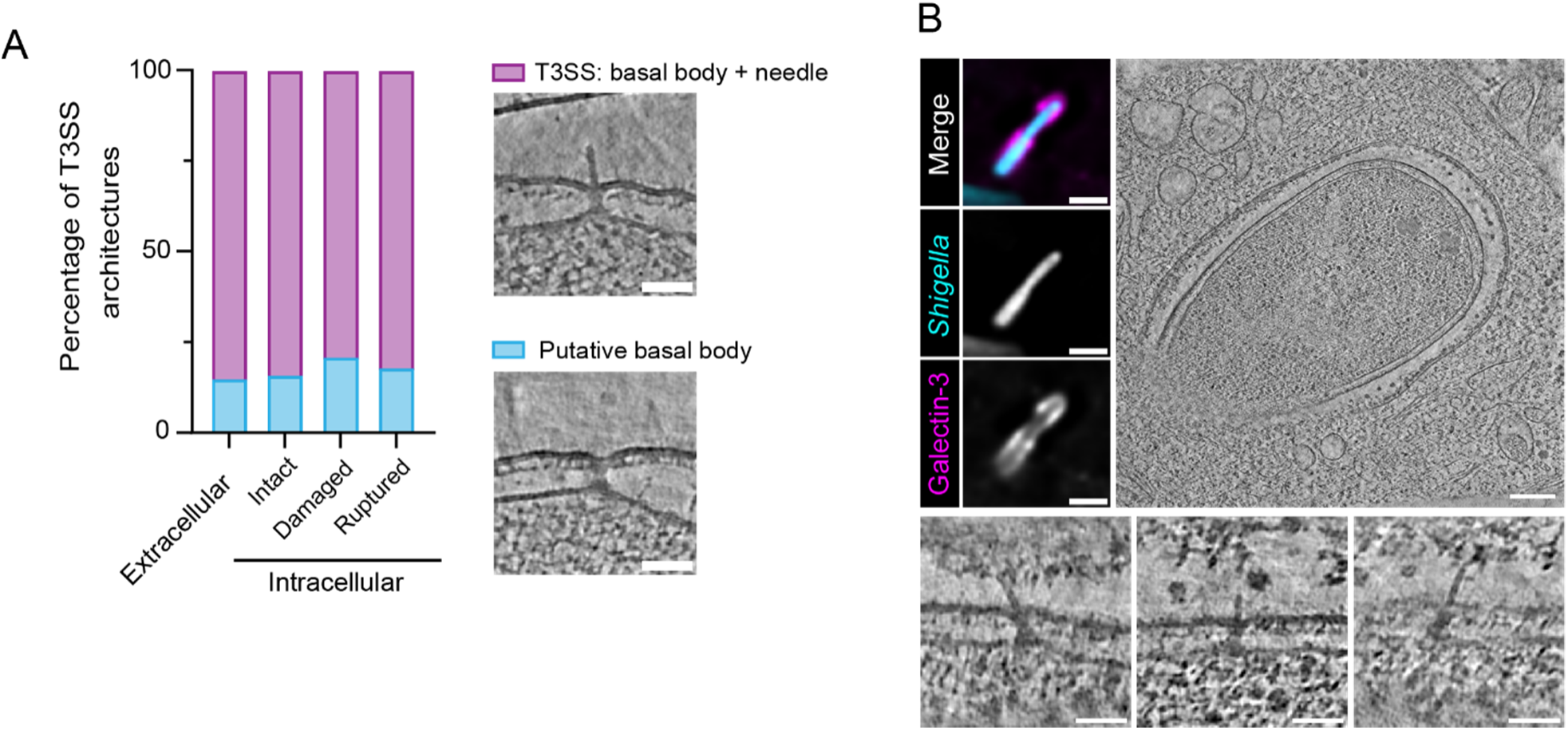
T3SS assemblies along *Shigella* infection stages. (A) Diversity of T3SS architectures might reflect different assembly states. Contingency graph of the percentage of T3SSs with a membrane-spanning basal body and protruding needle (total n=122) or with only a basal body without needles (total n=25) plotted according to the infection stage. Representative examples of putative T3SSs assembly states are shown. Scale bars are 50 nm. (B) T3SS needles are exposed to the cytosol after vacuole rupture and disassembly. Cryo-fLM, (scale bars 2 µm) and corresponding tomogram slice (scale bar 200 nm) of *Shigella* with ruptured vacuole displaying T3SSs with needles exposed to the cytosol after vacuole rupture (insets, scale bars 50 nm).

**Figure S4:**
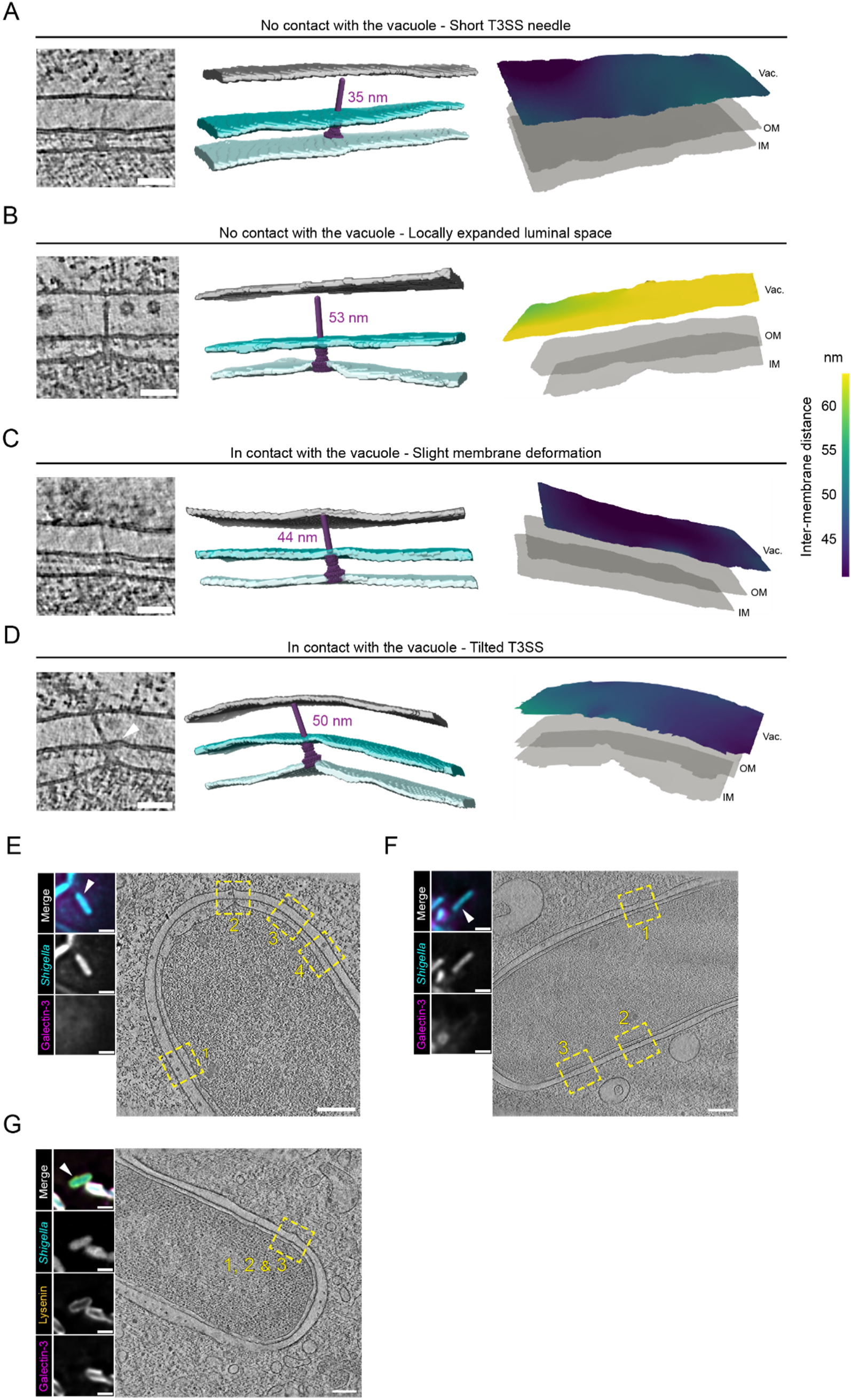
T3SS/vacuole contact depends on both T3SS length and available luminal space. (A-D) Tomogram slices of the T3SS/vacuole zones before *Shigella* membrane rupture. White arrowhead points to bacterial membranes deformations. Scale bars 50nm. Corresponding 3D rendering of vacuole and bacteria membranes with *Shigella* T3SS 3D map (EMD-15700) fitted. Galectin-3 negative vacuole (grey), bacteria OM (dark blue) and IM (light blue). Representative membrane surface reconstruction vacuole membranes coloured by relative distance to the bacteria outer membrane. Bacteria membranes are shown in light grey. Vac.: Vacuole, OM: Outer membrane, IM: Inner membrane.

(A and B) T3SSs do not establish contact with the vacuole if they are short (A) or if the vacuole is locally relaxed (B).
(C) T3SS with needle length correlating with the available vacuole space contact the vacuole without deforming it.
(D) T3SS with long needle contacting the vacuole membrane adopts tilted insertion within bacterial membranes. (E-F) Correlations of the tomogram insets shown in Figures 4 and S4. See also Supplementary Video S3. Slice through tomograms (scale bars 200 nm) and corresponding cryo-fLM images (scale bars 2 µm) of *Shigella* (white arrowhead) before vacuole rupture. Inset correspondence:

(E) T3SS 1: Figure S4B, 2: Figure 4B, 3: Figure S4D, 4: Figure 4A.
(F) T3SS 1: Figure 4C, 2: Figure S4D, 3: Figure S4A.
(F) T3SSs 1, 2 and 3: Figure 4D.

## List of supplementary videos

**Video S1. Time-lapse microscopy monitoring of Lysenin and Galectin-3 recruitment at the *Shigella* and *E. coli* mT3*_Shigella_* vacuole, related to Figure 1B and C**

*Shigella* and *E. coli* mT3*_Shigella_* infection of HeLa eGFP-Lysenin mOrange-Galectin-3 cells. Vacuolar damage characterized by Lysenin recruitment at the vacuole (merge: yellow, greyscale: second panel) is observed shortly before vacuolar rupture (merge: magenta, greyscale: third panel) indicated by the recruitment of Galectin-3 at the vacuole and is followed by vacuole disassembly. Images were taken every minute and a maximum intensity z-projection is presented. The scale bar is 5 µm.

**Video S2. Tomograms and corresponding segmentations of the distinct stages of *Shigella* cytosolic access, related to Figure 2A-C**

Sequential slices through the tomograms shown in Figure 2A, B and C and segmentation showing *Shigella* entrapped either into an intact or damaged vacuole or surrounded by ruptured vacuole remnants. *Shigella* OM: Outer membrane (dark blue), IM: Inner membrane (light blue). Vacuoles; intact vacuole (grey), damaged vacuole (yellow) and ruptured vacuole (magenta). Scale bars are 200 nm.

**Video S3. Tomograms and corresponding segmentations of *Shigella* prior cytosolic access, related to Figure S4E-G**

Sequential slices through the tomograms of Figure S4E, F and G with corresponding segmentations showing *Shigella* in Galectin-3 negative vacuoles or in a damaged vacuole. *Shigella* OM: Outer membrane (dark blue), IM: Inner membrane (light blue), Galectin-3 negative vacuole (grey) and damaged vacuole (yellow). Scale bars are 200 nm.

**Video S4. T3SS punctures host endomembrane, related to Figure 4C**

Sequential slices through a zone of tomogram showing a T3SS puncturing the vacuolar membrane. Scale bar is 50 nm.

